# Genetic variations in foraging habits and their developmental noise in *Drosophila*

**DOI:** 10.1101/2023.07.28.550901

**Authors:** Kaiya Hamamichi, Yuma Takahashi

## Abstract

Investigating the causes and consequences of niche partitioning in populations is a significant goal in ecology and evolutionary biology. Studies have examined genetic and environmentally induced variations in resource utility and their ecological implications. However, few have explored developmental noise or instability as a factor contributing to variation in resource utility. Here, we studied genetic variation, and developmental noise in foraging traits of *Drosophila lutescens*, a wild fruit fly. Using 70 iso-female lines from a single population, we observed two foraging behavior traits - locomotive activity and resource preferences - in an experimental “8”-shaped arena with two different fruit juices in each chamber. The mean locomotive speed and relative preference for orange juice over grape juice varied significantly among iso-female lines, indicating genetic variation in foraging behavior. Additionally, the degree of variation within iso-female lines also varied, showing relatively higher heritability. While the locomotive speed and resource preferences of each line did not correlate with each other, the strength of variation within iso-female lines for locomotive speed showed a significant correlation with that for resource preferences. This suggests that the degree of developmental noise in both locomotive activity and resource preferences is governed by a shared genetic basis. Consequently, developmental noise can contribute to increased phenotypic variation in resource utility within a population and may evolve through natural selection.

## Introduction

Niche and resource partitioning play a crucial role in ecology, enhancing community, and ecosystem functioning, stability, and productivity (1–4). Niche partitioning effectively reduces potential competition among individuals (5), and niche width reflects resource abundance along the resource axis, the pattern of intraspecific competition, and elements of natural selection (6). In recent decades, the ecological function of intraspecific resource partitioning has garnered increasing attention, with studies demonstrating the influence of niche partitioning, even within a single species, on population dynamics (7–10). Therefore, understanding resource and niche partitioning within a population is essential for comprehending population dynamics (11–14).

Phenotypic variation between individuals can arise from three causative factors: genetic variation, environmental variation (phenotypic plasticity), and stochastic developmental noise (15). Developmental noise, characterized by random fluctuations during development, contributes to variation among individuals sharing the same genetic and environmental background (16). The study of genetic and environmental variation in the context of intraspecific resource partitioning has been well-documented (17–21). For example, empirical laboratory experiments have provided evidence of a causal link between genetic differences and variations in resource preferences (22). Additionally, it has been established that the dietary preferences of the tobacco hornworm *Manduca sexta* are subject to plasticity and determined based on the previously consumed plant (23,24). However, few studies have explored developmental noise in foraging behavior and resource use within populations. In *Drosophila*, developmental noise in neuroplasticity has been linked to individual behavioral differences (25), indicating its potential influence on various traits, including foraging habits, and resource preferences. To comprehensively understand the mechanisms underlying intraspecific resource partitioning, both genetic variation, and developmental noise contributions must be thoroughly investigated.

This study aims to demonstrate the impact of genetic variation and developmental noise on variation in foraging habits in the wild fruit fly *Drosophila lutescens*. First, we established iso-female lines using females from a single natural population and evaluated genetic variation and developmental noise in two essential foraging traits, namely locomotive activity, and resource preferences, which shape resource partitioning. Subsequently, we estimated the heritability of each trait and its developmental noise, followed by an examination of the genetic correlation between the degree of developmental noise for each trait.

## Materials and Methods

### Sampling and study species

*Drosophila lutescens*, a member of the *melanogaster* group within the subgenus *Sophophora*, is considered a generalist species (26). In our study, To establish the iso-female lines, *D. lutescens* females were collected from the campus of Chiba University (35°37′34″ N, 140°6′9″ E) between January and April, as well as October and November, 2020. Each collected female was cultured in a plastic vial (φ30 mm and 100 mm in height) with a nutritive medium following the protocol of Fitzpatrick et al. (27) to establish iso-female lines. The composition of the medium was as follows: 500 ml of H_2_O, 50 g of sucrose, 50 g of active yeast, 6.5 g of agar, 5.36 g of KNaC_4_H_4_O_6_·4H_2_O, 0.5 g of KH_2_PO_4_, 0.25 g of NaCl, 0.25 g of MgCl_2_, 0.25 g of CaCl_2_, and 0.35 g of Fe_2_(SO_4_)·6.9H_2_O. Throughout the study, we used the same medium and vials for maintaining the lines. To ensure age homogeneity, individuals aged 1–7 days (postemergence) were used in all experiments, which were conducted in 2021–2022 using 70 established iso-female lines.

### Quantification of variation in foraging habits and heritability

To quantify foraging habits, an experimental arena with two confined chambers and a connecting passageway was used (Fig. 1). Each chamber contained either orange or grape juice as resources, with their positions in a concavity at the center of the chamber randomized for each measurement. Five females per iso-female line were introduced into the arena after being anesthetized using CO_2_. The flies were placed at the center of the passageway to minimize any initial position bias. Subsequently, the arenas were covered with slide glass, placed in an incubator (25°C), and recorded via a camera for 66 min. To mitigate the effects of anesthesia, we extracted coordinate data through automated tracking (Flytracker in MATLAB) from the last 44 min of the video. Five to eight replicates were used per line.

**Figure 1.**
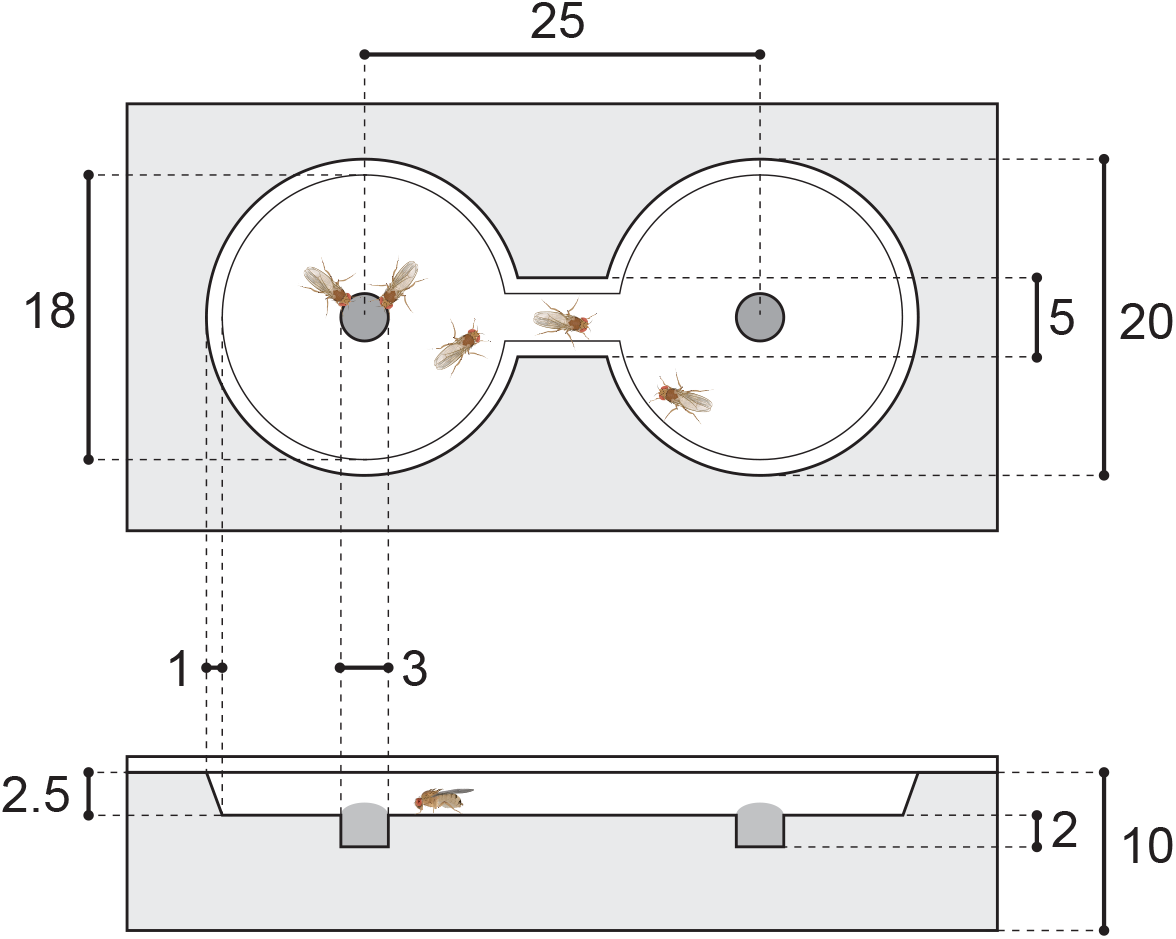
Anterior (upper) and transverse (lower) sections of the experimental arena. Black circles represent resources, whereas white regions indicate the area where individuals can walk. Unit of length: mm.

Resource preferences were estimated using two preference indices (*PI*s), calculated based on the time spent in the orange feeding chamber, the grape feeding chamber, and the passage. Two *PI*s were calculated as follows: *PI*_1_ = (*O* – *G*)/(*O* + *G* + *P*) and *PI*_2_ = log_10_(*O*/*G*), where *O, G*, and *P* represent the orange feeding chamber, grape feeding chamber, and passage, respectively. A higher *PI* value indicated a preference for orange, whereas a lower value indicated a preference for grape. A *PI* value of 0 indicated no preference.

To assess foraging activity, locomotive activity was quantified by calculating the average locomotive speed while searching. Data were excluded if the locomotive speed was <0.01 mm/s, which indicated that the fly had reached the food (3.5 mm from the center of the food). The locomotive activity was measured as the distance traveled every ∼0.67 s.

Phenotypic variation within each line, considered as developmental noise, was determined using the standard deviation (SD) for resource preferences and the coefficient of variation (%CV) for locomotive activity. The %CV for locomotive activity was calculated as the ratio of the SD to mean locomotive speed as follows: SD/M × 100, where M represents the mean.

The heritability of each trait and its developmental noise were evaluated using ANOVA, with each trait assigned as an objective variable and the lines assigned as explanatory variables. For developmental noise, fluctuations per arena (per five individuals) were used for heritability evaluation, as it cannot be assessed on an individual basis.

### Statistical analyses

All statistical analyses were conducted using R software (version 4.2.2). The R package *R*.*matlab* was used to read the coordinate data obtained from video tracking. To examine genetic variations among lines for each trait, generalized linear models were applied, assuming Gamma, quasibinomial, and Gaussian error distributions for locomotive speed, *PI*_*1*,_ and *PI*_*2*_, respectively. Pearson’s correlation test was used to examine correlations between locomotive speed and *PI* (*PI*_1_ or *PI*_2_) and between the %CV of locomotive speed and SD of *PI* (*PI*_1_ or *PI*_2_).

## Results

Preference for orange was predominant in most lines, with only a few lines exhibiting a preference for grape (Fig. 2a; S1 Figure). The variations in preferences were significant among the lines (*PI*_1_: *df* = 69, χ^2^ = 147.9, *P* < 0.001; *PI*_2_: *df* = 69, χ^2^ = 114.1, *P* < 0.001). The heritability of preferences, calculated through *PI*_1_, and PI2, was *H*^2^ = 5.1% and *H*^2^ = 4.0%, respectively. The degree of intraline variation for resource preferences varied among the lines. For *PI*_1_, the heritability of the degree of individual variation within lines was *H*^2^ = 16.7%, whereas that for *PI*_2_ was *H*^2^ = 22.6%.

**Figure 2.**
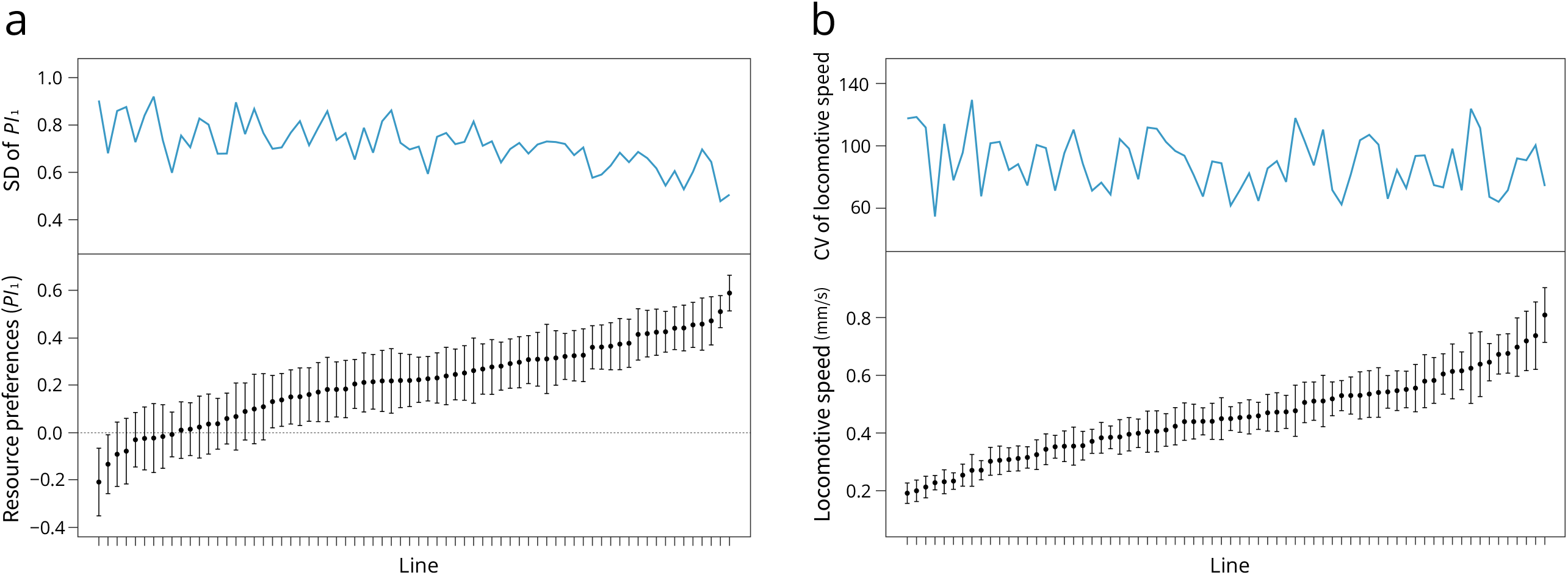
Mean (lower) and intraline (upper) variation for *PI*_1_ (a) and locomotive speed (b). Lines are arranged in ascending order based on their respective means, and error bars represent standard errors.

Locomotive activity also exhibited significant variation among the lines, with the largest, and smallest lines exhibiting locomotive speeds of 0.8 and 0.2 mm/s, respectively (*df* = 69, χ^2^ = 347.2, *P* < 0.001; Fig. 2b). The heritability of locomotive activity was estimated as *H*^2^ = 9.9%. The degree of individual variation (%CV) ranged from 50% in the smallest line to 130% in the largest line. The heritability of intraline variation for locomotive activity was *H*^2^ = 10.0%.

Although no significant correlation was observed between mean resource preferences and mean locomotive speed, a significant correlation was found between the intraline variation of resource preferences and that of locomotive activity (Fig. 3a, b; S2 Figure a, b). This suggests that lines with greater variation in locomotive activity also exhibit greater variation in resource preferences.

**Figure 3.**
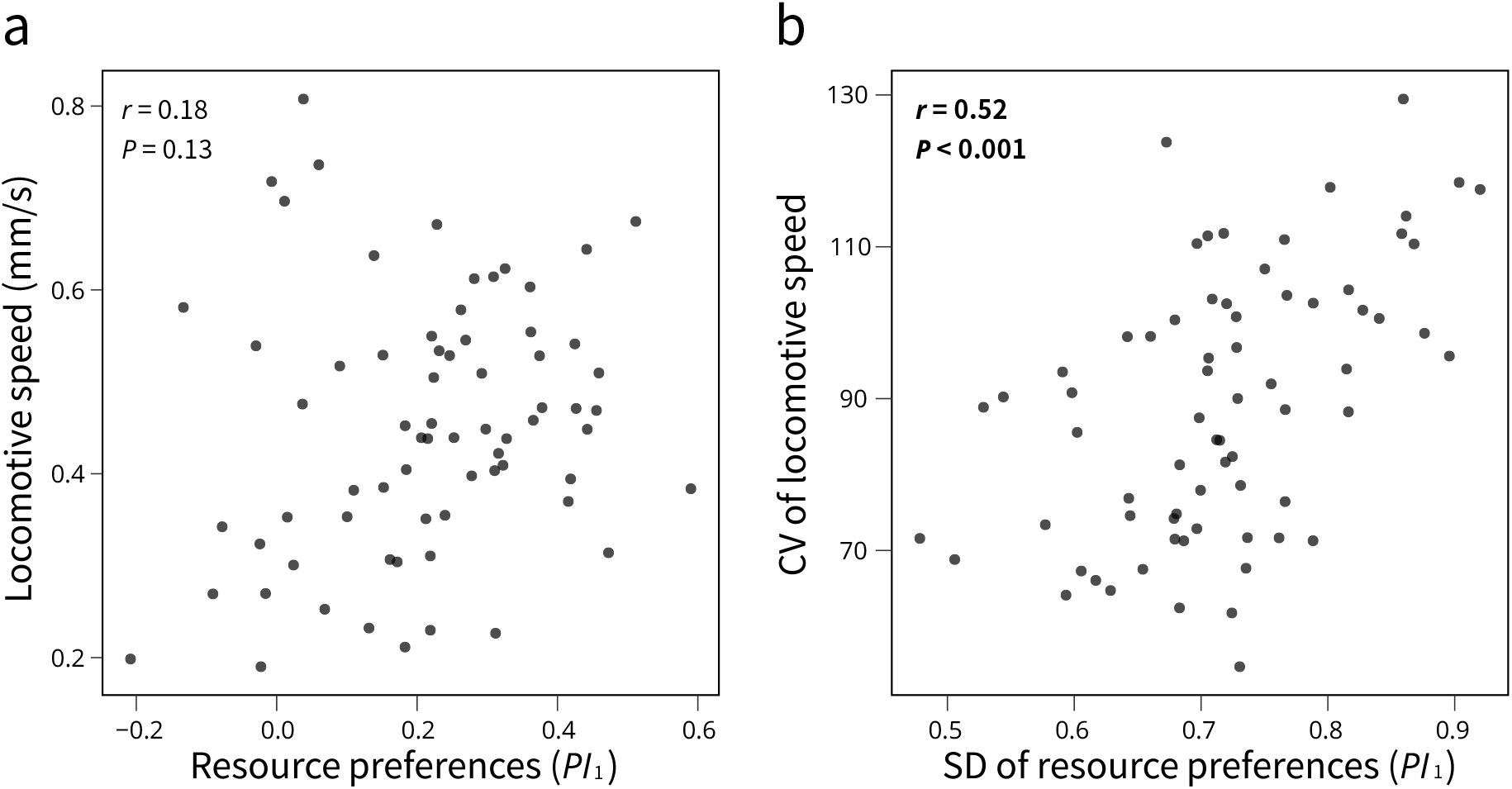
Correlations between mean locomotive speed and mean *PI*_1_ for each line (a), and correlations between intraline variation in locomotive speed and in *PI*_1_ for each line (b). Each panel displays correlation coefficients and p-values determined using Pearson’s correlation test.

## Discussion

Despite the ecological significance of intraspecific niche partitioning (7–10), limited research has focused on investigating the impact of developmental noise on such partitioning. In our study, we uncovered variations resulting from both genetic diversity and developmental noise in foraging habits within the population. Notably, we found that the degree of developmental noise was heritable, indicating its potential role in contributing to increased phenotypic variation in resource utility among individuals within the population. Consequently, developmental noise emerges as a crucial determinant capable of influencing ecological dynamics.

The intraline variation quantified in the present study may potentially include measurement errors. If the quantified intraline variation resulted from measurement error, the heritability of noise would be zero, and the noise of the two independent traits should not exhibit any correlation with each other. However, our findings indicate that the strength of intraline variation was heritable, exceeding 10%, and we observed a significant positive correlation in the intraline variation of the two traits, implying that the noise observed in locomotive activity and resource preferences is not solely due to measurement error but rather reflects developmental noise in traits. Moreover, the significant positive correlation in the developmental noise of the two traits suggests that a shared genetic basis governs the degree of developmental noise in both locomotive activity and resource preferences. Certain genetic factors may contribute to the occurrence of developmental noise across multiple traits, as observed in plants, and *Drosophila*, with the protein Hsp90 known to govern the degree of developmental noise in multiple phenotypes (28,29). The evolution of genes controlling the expression of such proteins may influence the magnitude of diversity resulting from developmental noise. Further studies are needed to explore the relationship between developmental noise and fitness (30). Note that, at this time, we cannot rule out the possibility that intraline variation quantified in our study partly reflected the degree of genetic variation within a line and the effects of plastic variation derived from environmental heterogeneity shaped by line-specific ecological engineering, though we strived as much possible as to eliminate the influence of genetic and plastic factors on variation within a line.

Resource preferences and locomotive activity may directly or indirectly lead to intraspecific niche partitioning, which has crucial ecological implications, including avoiding competition (14,31). The occurrence of intraspecific niche partitioning has been reported in various organisms (32,33). Alongside genetic variation, phenotypic plasticity, and developmental noise could contribute to generating variation in foraging habits within a population, thereby mitigating resource competition among individuals. However, the present study did not determine the relative contribution of genetic variation and developmental noise in generating this variation. Consequently, further experimental studies, using iso-female lines with different degrees of developmental noise for foraging habits, are required to clarify the contribution of developmental noise in generating variation within a natural population and its effects on population dynamics.

## Acknowledgments

This work was supported by research grants JSPS KAKENHI Grant Numbers 22H05646 and 23H03840, Toyota Foundation, and Sumitomo Foundation to Y.T.

## Data Accessibility

All raw data are available in Figshare (doi: 10.6084/m9.figshare.23747442).

## Author Contributions

K.H. and Y.T. designed the experiments. K.H. performed the experiments and conducted the data analysis. K.H. drafted the manuscript. K.H. and Y.T. edited the manuscript.

