## Supplementary material for "Genetic variations in foraging habits and their developmental noise in *Drosophila*": SI

### Supporting Information

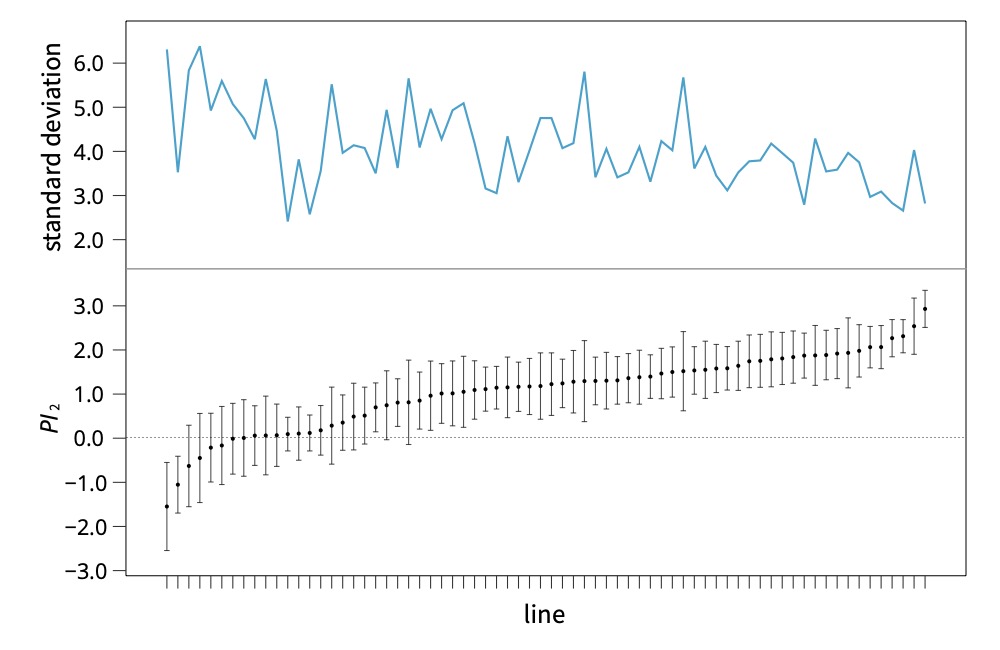

**S1 figure.** The mean (lower) and variation (upper) in *PI*_2_ in each line. The lines are sorted in ascending order of mean *PI*_2_, and error bars indicate standard errors.

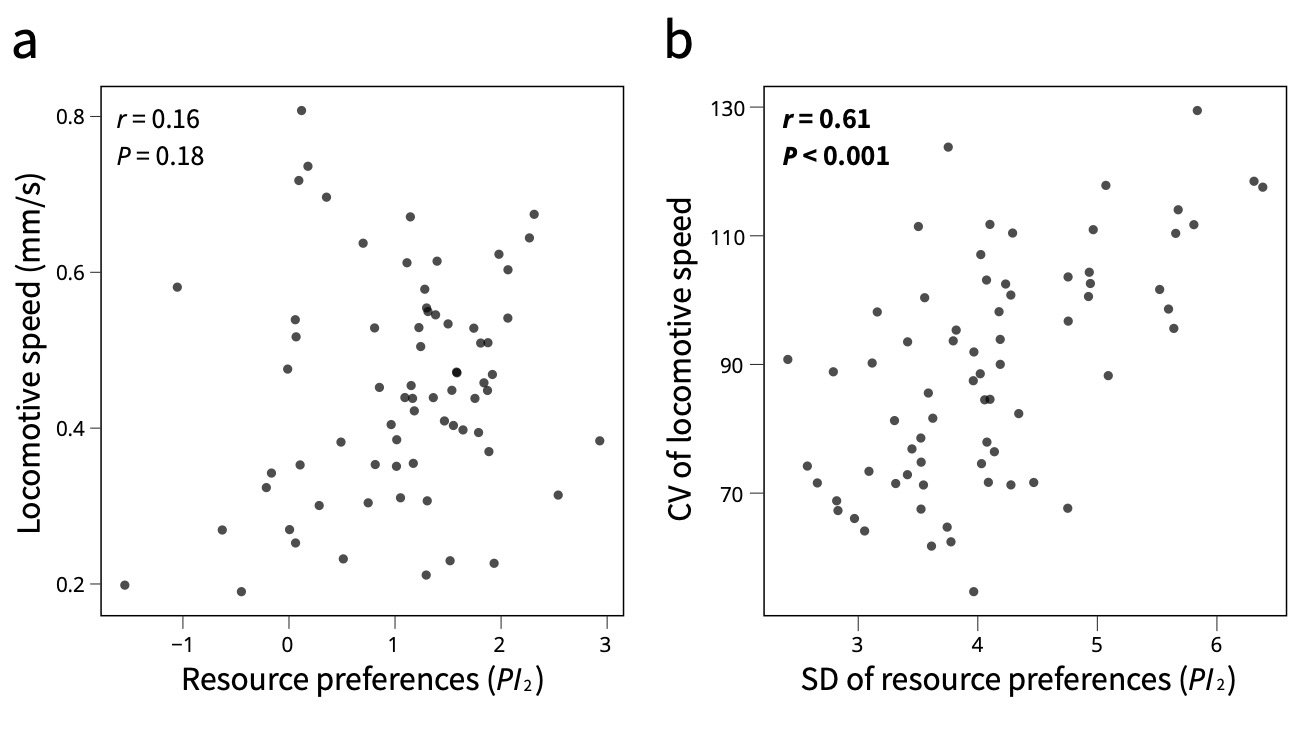
**S2 figure.** The correlation between the mean locomotive speed and the mean *PI*_2_ for each line (a), and the correlation between intra-line variation in locomotive speed and that in *PI*_2_ (b). The correlation coefficient and p-value examined by Pearson’s correlation test are shown in each panel.
